# Whole Genome Assembly of the Atlantic Bluefin Tuna *Thunnus thynnus,* with Phased Haplotypes

**DOI:** 10.1101/2025.11.18.689121

**Authors:** Barbara A. Block, Shaili Johri, André Machado, Adam Ciezerak, Filipe Castro, Paul Peluso, Greg Concepcion, Tal Eli-Zaquin, Laia Juliana Del Rocio

## Abstract

Atlantic bluefin tuna (*Thunnus thynnus*) are highly migratory fish, the largest tuna in the genus *Thunnus*. These fish possess a unique suite of traits, including regional endothermy, in their swimming muscle, viscera, brain and eyes that elevates tissue temperatures significantly above ambient water temperatures. Bluefin tuna also have an efficient ‘thunniform’ locomotory style, involving a fusiform low drag external shape and a novel musculoskeletal system that enhances elastic energy storage. Together the unique swimming form and warm muscle temperatures enable this species to move across large expanses of the ocean. Atlantic bluefin tuna are targeted by high-value international fisheries of over fifty nations. Populations have recovered in recent years and the species is now actively farmed extensively within the Mediterranean Sea and eastern Atlantic. To date, fisheries management and evolutionary studies have been hampered by a lack of genomic resources. Here, we report a novel *de novo* chromosome-level phased genome assembly for the Atlantic bluefin tuna obtained using PacBio HiFi reads and Dovetail Omni-C scaffolding. This assembly was scaffolded into 24 chromosomes containing 98% of the genes in the BUSCO reference dataset, demonstrating a high level of contiguity with a contig N50 of 26.54 and assembly size of ∼800Mbp. We compare a diploid-resolved (2N) genome assembly of Atlantic bluefin tuna with a haplotype-collapsed (1N) genome assembly from a Pacific bluefin tuna caught in the eastern Pacific off California. We show the two species have a 98.47% sequence identity. These high-quality genomes for northern bluefin tunas provide the genomic tools necessary for improving phylogenomic and evolutionary physiology analyses of this lineage and are important for potential functional assays for population biology of these commercially important species.

## Introduction

The Atlantic bluefin tuna (ABFT), *Thunnus thynnus,* and its sister species the Pacific bluefin tuna (PBFT), *Thunnus orientalis,* are northern bluefins, the largest extant members of the *Thunnus* genus (Shimose & Farley, 2015). These iconic fish are ecologically and economically significant, highly sought after by commercial and recreational fishers for their remarkable endurance during pole and line fishing. Over the past half century, significant population declines have led the International Union for Conservation of Nature (IUCN) to classify ABFT as vulnerable and PBFT as endangered (Collette et al., 2021; Juan-Jordá et al., 2011). Importantly, reductions in international fishing pressure across both the Atlantic and Pacific Oceans have facilitated a steady recovery in their populations, particularly in the eastern Atlantic and the Mediterranean Sea.

The large size, power, and speed of northern bluefin tuna pose challenges for ecological and physiological studies, but recent advancements in biologging tags have significantly enhanced our understanding of their ecology, behavior, and physiology over the past two decades (B. Block et al., 1998; B. A. Block et al., 2019; B. Block & Stevens, 2001). ABFT and PBFT occupy a large home range in their respective oceans, from sub-tropical to cool sub polar waters, as far north as Iceland, the Faroe Islands and Norway (B. A. Block et al., 2019; MacKenzie et al., 2019). Recent catches of ABFT in northern regions indicate a significant return to their historical range after a prolonged absence, particularly off the Faroe Islands, Scotland, and Iceland (Makenzie et al. 2014, Horton et al. 2021). Northern bluefin tunas have dive rapidly from the warm surface waters to cooler waters at depth below the thermocline, often feeding on mesopelagic animals at 300-600m and in some cases well over 1000m deep (Pagniello et al., 2023; Walli et al., 2009). The evolution of regional endothermy in bluefin tunas, whereby elevated temperatures are maintained in deep red swimming muscle, cranial and visceral tissues by the presence of retia mirabilia enables the fish to maintain body temperature significantly above ambient (B. A. Block & Finnerty, 1994). Endothermy is associated with high aerobic capacity in the tissues, an increase of slow-twitch muscle volume and increased capacity for excitation-contraction coupling in the heart (Blank et al., 2007; Shiels et al., 2011), that improves cardiac function in the cold. Conservation of metabolic heat increases muscle power output, endurance performance, and the speed of digestion (Watanabe et al., 2015; Whitlock et al., 2015). Warming of the brain and eyes has been shown to improve vision in other species with cranial endothermy (Fritsches et al., 2005).

The nothern bluefin range extensively across the Pacific and Atlantic basins, respectively. The southern bluefin tuna, *T. maccoyii*, has a smaller body size but similarly can also tolerate cold temerate waters and migrates to high-latitude feeding areas (Arrizabalaga et al., 2015; Shimose & Farley, 2015). Despite their similarities, phylotranscriptomic and RAD-seq based studieshave indicated that the three bluefin species are paraphyletic. The southern bluefin is more closely related to warm-water *Thunnus* species such as bigeye tunas, whereas the two northern species are sister taxa to each other and only recently diverged (Ciezarek et al., 2019; Díaz-Arce et al., 2016). ABFT and PBFT have distinctly different stock structures. Tagging, genetics and fisheries catch date indicate there is a single PBFT stock in the Pacific ocean (Nakatsuka, 2020), with their distributions primarily in the north although they undertake a post spawning migration in the western Pacific into New Zealand waters (Fujioka et al., 2015). By contrast, there are at least three distinct populations of ABFT, which mix on shared foraging grounds in the north Atlantic, but have unique spawning grounds in the Gulf of Mexico, the Mediterranean Sea as well as the slope waters of the mid-Atlantic blight (Pagniello et al., 2023; Richardson et al., 2016; Rooker et al., 2008).

Understanding the evolutionary dynamics and functional diversification of gene families is essential for elucidating the adaptive strategies of bluefin tunas and their relatives within the Scombridae family. By examining orthologous relationships among selected teleost species—including the recently annotated genomes of *Thunnus orientalis*, *Thunnus thynnus*, and *Scomber scombrus*—we can gain insights into the genetic bases of traits that enhance survival in challenging environments, such as thermal tolerance and hypoxia resistance (Emms & Kelly, 2019). Analyzing gene family evolution will provide a critical foundation for conservation genomics efforts aimed at preserving bluefin tuna populations amidst environmental changes and fishing pressures.

As bluefin tuna stocks may be influenced differentially by fisheries exploitation and climate change, it is important to build the genetic resources necessary to examine the evolutionary history and the physiological variation of different bluefin tuna populations. Conservation genomics offers the potential to better manage these populations, by enabling accurate population delineation and understanding patterns of diversity within and between populations, the extent any extent of gene flow or inbreeding, and the identification of adaptive mutations (Bernos et al., 2020; Supple & Shapiro, 2018). Currently, the impact of population declines, and trajectories of recovery in some populations as a result of recent conservation efforts on genetic diversity of the bluefin tunas is unknown. Genomics will also aid the growing interest in high-value aquaculture of bluefin tunas (Klinger & Mendoza, 2019), by allowing identification of genes for important traits such as growth. A necessary first step for selective breeding is the generation of a high-quality reference genome. Recent studies have published *de novo* and draft genomes for PBFT (Nakamura et al., 2013; Suda et al., 2019), southern bluefin tuna (Zhao et al., 2022), and warm-water yellowfin tuna (Dimens et al., 2024). In this context, we present a chromosome-level assembly for ABFT and compare it to a collapsed assembly of PBFT from the eastern Pacific Ocean.

Advancements in sequencing technologies have greatly enhanced genomic research capabilities, with the HiFi methodology from Pacific Biosciences standing out for its ability to provide read lengths of 10 kb to 20 kb and high accuracy rates exceeding 99.5% at the single-molecule level (Wenger et al., 2019). These HiFi reads have shown considerable effectiveness across various genomic applications, including small variant detection, structural variant identification, and de novo assembly of complex genomes. This is exemplified by their performance in characterizing the human male genome HG002/NA24385, part of the Genome in a Bottle (GIAB) Consortium (Zook et al., 2016, 2019). The technology has proven instrumental in addressing challenges such as closing gaps associated with segmental duplications in genomes (Nurk et al., 2020; Vollger et al., 2020) and resolving haplotypes without relying on pedigree information (Garg et al., 2021). Given the complexities involved in studying deep ocean fish genomes, where pedigree data may be sparse, the integration of both HiFi and Omni-C linked reads (Dovetail Genomics, Scotts Valley, CA, USA) is particularly beneficial.

The primary aim of this study is to develop a high-quality reference genome for T. thynnus, facilitating a deeper understanding of its evolutionary dynamics and the functional diversification of gene families. Through the analysis of orthologous relationships among key teleost species, including the annotated genomes of *Thunnus orientalis*, *Thunnus thynnus*, and *Scomber scombrus*, we seek to uncover significant insights into the genetic foundations of adaptive traits such as thermal tolerance and hypoxia resistance. This work is also intended to lay crucial groundwork for conservation genomics initiatives that focus on the preservation of bluefin tuna populations in response to environmental changes and fishing pressures. Gaining a better understanding of gene family evolution is essential not only for the biology of bluefin tunas but also for guiding effective management strategies aimed at ensuring the sustainability of these vital species within their ecosystems.

## Materials and Methods

### Genomic DNA Isolation

Genomic DNA (gDNA) was isolated from an Atlantic bluefin tuna male with 220cm in length and 108kg in weight, which was caught in the waters north of Cape Hatteras, North Carolina in the USA. The gDNA was isolated using a Qiagen Genomic Tip 500/G DNA extraction kit following the manufacture’s protocol. A total of 400 mg of tissue were used for the DNA extractions of the Atlantic bluefin tuna. The sample was obtained from the ventricle of a tuna heart that was collected in February of 2019 from a commercially caught fish off the coast of Hatteras, North Carolina. The heart was packed on ice, shipped overnight to the lab without freezing, and the DNA was isolated within 24 hours. DNA concentration was quantified with the Qubit High Sensitivity Kit (Invitrogen, USA) and DNA purity was quantified with a Nanodrop spectrophotometer (ThermoFisher, USA). Samples for Pacific bluefin tuna were obtained from a fish caught in the California Current and isolation protocol was similar.

### HiFi Library Preparation

Genomic DNA was converted into a SMRTbell™ library as previously described (Wenger et al., 2019) but with a few modifications to generate slightly larger inserts. Specifically, gDNA was sheared using the MegaruptorR from Diagenode with the 20kb shearing protocol using a long hydropore cartridge. Prior to library preparation, the size distribution of the sheared DNA was characterized on the Agilent Femto Pulse System. Sequencing libraries were constructed from sheared genomic DNA using the SMRTbell™ Express Template Prep Kit v 2.0 and subsequent treatment with the Enzyme Clean Up Kit (Pacific Biosciences Ref. No. 101-843-100). In order to tighten the size distribution of the SMRTbell™ library, library was size fractionated using SageELF System from Sage Science. Approximately 8 µg of SMRTbell™ Library, prepared with loading solution/Marker40. After which, the sample was loaded onto a 0.75% agarose 10kb-40kb gel cassette and size fractionated using a run a target size of 7000bp set for elution well 12. A total of 8µg was fractionated on two cassettes. Fractions having the desired size distribution ranges were identified on the Agilent Femto Pulse System. Fractions centered at 20kb were pooled to generate the final size-selected library used in the sequencing runs.

### HiFi reads sequencing

Sequencing reactions were performed on the PacBio Sequel II System with the Sequel Sequencing Kit 2.0 chemistry. The kit uses a circular consensus sequencing (CCS) mode which provides >99% single-molecule read accuracy allowing resolution of single nucleotide as well as structural variants. The samples were pre-extended without exposure to illumination for 4 hours to enable the polymerase enzymes to transition into the highly progressive strand-displacing state and sequencing data was collected for 30 hours to ensure maximal yield of high-quality CCS reads. CCS reads were generated from the data using the SMRT Link (v.8.0).

### Omni-C reads sequencing

In order to obtain chromosomal level scaffolds proximity ligation libraries were generated using the Omni-C kit and protocol (Dovetail genomics). Briefly, for each library, 20mg of heart tissue was fixed with formaldehyde then digested *in situ* with DNAse I. Digested ends were repaired and capped with a biotin-containing bridge sequence. The bridge sequences were proximally ligated, and the resulting DNA was used to generate Illumina-compatible sequencing libraries, using the NEBNext Ultra II DNA Library Prep kit. After adapter ligation, libraries were enriched for biotin-containing molecules prior to PCR amplification. The resulting libraries were sequenced to a depth of 100-200M reads on an Illumina NextSeq 550 platform.

### Genome assembly and scaffolding

Pacific Biosciences high-fidelity (HiFi) sequencing reads, combined with the hifiasm assembler, resulted in genome assemblies at nearly complete haplotype resolution. The various assemblies of Atlantic bluefin tuna were compared with a publicly available Pacific bluefin tuna assembly (Suda et al., 2019). Additionally, a new haplotype-collapsed Pacific bluefin tuna genome was generated from a fish captured in the California Current, using PacBio HiFi reads and scaffolded with Hi-C data.

### Validating haplotype phasing accuracy

Because there was no parental information available for the sequenced ABFT sample, phasing accuracy could not be determined by direct comparison, as in the case of HG002 genome done at the same time. Instead, haplotypes separation was validated based on k-mer abundance in each haplotype using the k-mer analyses tool (KAT) (Mapleson et al., 2017).

### Haplotype comparison for sequence and structural variation

Variant quality, depth, type and frequency, along with genotype quality, type and frequency were determined using DeepVariant (https://github.com/google/deepvariant/blob/r0.10/docs).

### Synteny

To investigate chromosome structure and genomic synteny, the phased haplotype 1 chromosomal genome of Atlantic bluefin tuna (T. thunnus: GCA_043381625.1) was aligned with the phased haplotype 2 genome (T. thunnus: GCA_043381665.1), as well as with the previously published assembly of Pacific bluefin tuna (T. orientalis: GCA_009176245.1). For this analysis, all chromosomal scaffolds from both genomes were aligned using scripts from https://github.com/mcfrith/last-genome-alignments and LASTv921 (Kiełbasa et al., 2011). The process began with creating an index for each genome using the “lastdb” function of LAST with the options “-P0,” “-uNEAR,” and “-R01.” Substitutions and gap frequencies were then determined using the “last-train” algorithm with the options “-P0,” “--revsym,” “--matsym,” “--gapsym,” “-E0.05,” and “-C2.” Many-to-one alignments were generated with “lastal” using the options “-m50,” “-E0.05,” and “-C2,” followed by the application of the “last-split” algorithm with the option “-m1.” These many-to-one alignments were subsequently filtered to create one-to-one alignments utilizing “maf-swap” and “last-split” with the option “-m1.” Simple sequence alignments were removed using “last-postmask,” and the final output was converted to tabular format via “maf-convert -n tab.” Alignments were visualized using the CIRCA software (http://omgenomics.com/circa), and mismapping statistics were calculated. Alignments shorter than 10 kbp and those with an error probability greater than 1 × 10−5 were filtered out.

### Variant Classification

Variants were categorized according to their specific characteristics. Insertions were identifiedas instances where the alternative allele (ALT) was longer than the reference allele (REF), whereas deletions were characterized by the ALT being shorter than the REF. Single nucleotide polymorphisms (SNPs) were defined as variations affecting a single base within both ALT and REF. Biallelic variants contained one alternate allele, while multiallelic variants had multiple alternate alleles. Each variant was counted once; multiallelic variants were labeled as “Multiallelic_Deletion/Insertion/SNP” if all alternate alleles fell under the same type (e.g., all insertions), or as “Multiallelic_Complex” if they included different types. Reference calls (RefCalls) matched the reference genome and, although not classified as variants, were included in the overall dataset. The distribution of sequencing depth was evaluated through a histogram, reflecting the depth values found in the dataset. This histogram included all entries, encompassing RefCalls, while those without a depth measurement were omitted from the analysis.

### Quality Score Assessment

Quality scores, representing the confidence in the reported alternative alleles, were derived from the dataset. Higher scores indicated a lower likelihood of miscalling a variant, and these scores were visualized in histograms across the complete dataset, including RefCalls.

### Genotype Quality Evaluation

Genotype quality scores were extracted from relevant data fields. It is crucial to recognize that a high quality score suggests confidence in detecting the variant, while the genotype quality score may be lower if there is uncertainty regarding whether the variant is heterozygous or homozygous. Genotype quality was calculated on the Phred scale based on the formula -10* log10(probability of incorrect genotype assignment). Entries lacking genotype quality scores were excluded from this analysis.

### Variant Allele Frequency Distribution

The distribution of variant allele frequencies (VAFs) across different genotypes was depicted in histograms, with guiding lines illustrating expected VAFs for significant genotypes. For instance, heterozygous variants were expected to show a VAF near 0.5, indicating a balanced distribution of variant-supporting and reference-supporting reads. Reference calls typically exhibited a VAF above 0, as they would otherwise be designated as variants. Genotypes were categorized based on their descriptive values, with 0/1 and 0/2 classified as heterozygous (0/x) and 1/1 and 3/3 as homozygous (x/x). Instances of heterozygous genotypes with distinct alternate alleles, such as 1/2 or 3/5, were represented as (x/y).

### Biallelic Base Change Frequency

Counts of biallelic SNPs were recorded based on the frequency with which specific reference alleles transitioned to corresponding alternate alleles, as shown in the relevant charts.

Reference calls and multiallelic variants were excluded from this assessment. Additional insights into specific base changes can be referenced in the discussion of the Ti/Tv ratio.

### Biallelic Ti/Tv Ratio Analysis

The count of transitions (Ti) was defined as the number of biallelic SNPs resulting from changes between purines (A-G) or pyrimidines (C-T). Conversely, transversions (Tv) involved changes between different nucleotide types. A Ti/Tv ratio significantly exceeding one is regarded as indicative of a robust transition rate, reflecting the molecular structure of nucleotides. Pertinent literature provided context for interpreting this ratio. All biallelic SNPs were included in this analysis, with reference calls being excluded.

### Distribution of Biallelic Indel Sizes

The size distributions of biallelic insertions and deletions were illustrated using histograms, represented in both linear and logarithmic scales. Reference calls and multiallelic variants were omitted from these distributions. Structural variations and chromosomal alignments were analyzed employing the Progressive Mauve software (Darling et al., 2010).

### Repeat Masking and Genome Annotation

The genome annotation was performed for the ABFT chromosome-scale haplotype 1. Before proceeding with the gene structure prediction and functional annotation we identified and mask the ABFT repetitive elements. RepeatModeler v.2.0.1 (Flynn et al., 2020) software was initially used to create a *de novo* library of repeat elements in ABFT genome. Next, the *de novo* library in combination with libraries of Dfam_consensus-20170127 and RepBase-20181026 databases were used to soft mask ABFT genome in RepeatMasker v.4.0.7 (Tarailo-Graovac & Chen, 2009) software. The structural gene annotation was performed with both the MAKER v3.01.03 (Cantarel et al., 2008), and AUGUSTUS v2.5.5 (Stanke et al., 2006) pipeline as well as the BRAKER2 pipeline v2.1.6 (Brůna et al., 2021; Hoff et al., 2016, 2019) in parallel, using multiple sources of evidence - RNA-Seq of the same species ; proteins of multiple chordates and transcripts of Scombridae species (obtained from genome). The complete protein and RNASeqdatasets (CDD, Coils, GO, Gene3D, Hamap, InterPro, KEGG, MetaCyc, MobiDBLite, PIRSF, PRINTS, Pfam, ProSitePatterns, ProSiteProfiles, Reactome, SFLD, SMART, SUPERFAMILY, TIGRFAM ) were gathered from GeneBank database, aligned to the soft-masked ABFT genome with Hisat2 v.2.2.0 (D. Kim et al., 2015; H. Kim et al., 2019) with the default settings, and then the alignments sorted with SAMtools v.1.9 (Danecek et al., 2021).. While the protein data set included 130 Actinopterygii species and 16 other species spread across the Chordata phylum, the transcriptome datasets only included Scombridae species (*Thunnus albacares* and *Thunnus maccoyii*). The Scombridae transcriptomes were mapped to the ABFT genoma, with minimap2 v.2.24 (Li, 2018), (Paremeters: -ax splice:hq) software, and converted/sorted with SAMtools v.1.9 (Danecek et al., 2021). Finally, both RNASeq and transcriptome alignments were used as input to the MAKER pipeline and RNASeq, transcriptome alignments as well as proteomes used for the BRAKER2 pipeline (Paremeters:–etpmode; –softmasking; –UTR = off; –crf; –cores = 30), to predict gene structures. The BRAKER2 .gff file was then filtered by evidence, and all gene structures supported by at least one source of evidence (RNASeq or protein) collected for further analyses (BRAKER2 auxiliary scripts; selectSupportedSubsets.py). The post-processing stage was carried out by Another Gtf/Gff Analysis Toolkit (AGAT) v.0.8.0 tool (Dainat J, 2021). In AGAT the .gff3 file was uniformized and corrected for acronym names, overlap coordinate regions, start and stop codons and filtered for incomplete genes. The final set of proteins from the BRAKER2 pipeline were collected and functional annotated with InterProScan v.5.44.80 (Jones et al., 2014) tool and BLASTP searches. The BLASTP searches were performed with the DIAMOND v.2.0.13 (Buchfink et al., 2014) against RefSeq (O’Leary et al., 2016) and UniProtKB/SwissProt (The UniProt Consortium, 2021) DBs (Download at: 10/03/2022).

### Topology Weighting analysis

Recent phylogenetic studies have indicated that the southern bluefin tuna, *T. maccoyii,* is more closely related to the warmer water tunas than the two northern bluefin tuna, *T. orientalis* and *T. thynnus* (Ciezarek et al., 2019; Díaz-Arce et al., 2016). However, phylogenetic relationships vary across the genome, largely due to incomplete lineage sorting and potentially introgression (Maddison, 1997). Therefore, we carried out a topology weighting analysis to examine whether there are regions of the genome where the southern bluefin tuna is more closely related to the two northern bluefin tuna than the yellowfin tuna, and whether genes in these regions may explain the high-latitude adaptations of the three bluefin species.

To generate the dataset for this, a genome-wide alignment was generated using the novel Atlantic bluefin, Pacific bluefin and yellowfin tuna genomes published here, a southern bluefin tuna genome assembly published as part of the Darwin tree of life project. The alignment was generated using the reference-free aligner progressive cactus (J. Armstrong et al., 2020), and converted to maf format using mafTools (Mayakonda et al., 2018). These were then converted to fasta format using the galaxy toolkit (Ostrovsky et al., 2021), with one concatenated sequence for each of the assembled linkage groups, using both the PBFT and ABFT sequence as the reference separately. To run twisst, these fasta sequences were then converted to geno format, and phylogenetic trees were inferred in sliding windows of 200 SNPs, with 40 SNP overlap using iqtree (v2.3.0), using scripts adapted from genomics_general. TWISST was then run to carry out topology weighting analysis along the linkage groups, with results plotted in R. The relative frequencies of two topologies in particular were compared: the ‘species-tree’ topology (Mackerel,((SBFT,Yellowfin),(ABFT,PBFT)));, and a ‘bluefin-tree’ topology (Mackerel,(Yellowfin,(SBFT,(ABFT,PBFT))));

### Characterization of Orthologous Relationships and Gene Family Evolution

To characterize orthologous relationships and investigate gene family evolution across teleosts, sequences with a minimum length of 30 amino acids were clustered into hierarchical orthogroups (HOGs) using OrthoFinder version 2.5.5, with human sequences included as an outgroup (Emms & Kelly, 2019). Teleost species were selected based on their representation of significant lineage diversification leading to the Scombridae order, including the final annotation sets for *Thunnus orientalis*, *Thunnus thynnus*, and Scomber scombrus.

HOGs were categorized into three distinct groups: Unclustered HOGs (single-sequence HOGs), Single-species HOGs (comprising paralogous genes from a single species), and Multispecies HOGs (represented by homologous genes from at least two species). Multispecies HOGs were further classified into Single-copy genes (lacking same-species paralogs) and Multi-copy genes (containing same-species paralogs).

To determine the putative functional classes of gene families, sequence annotations for T. thynnus were conducted using InterPro version 5.65-97.0 (Jones et al., 2014; McDowall & Hunter, 2011) and eggnog-mapper version 2.1.12 (Cantalapiedra et al., 2021; Huerta-Cepas et al., 2019). Gene ontology (GO) term enrichment for selected genes in each categorization was assessed utilizing the topGO R package with the Fisher’s exact test, applying the “weight01” algorithm and establishing significance at p < 0.01 (Alexa & Rahnenführer, 2009).

### Divergence Time Estimation

A concatenated dataset comprising 1,690 single-copy orthologous HOGs, identified across the teleost lineage, was processed using trimAl version 1.4.1 (Capella-Gutiérrez et al., 2009) with the -gappyout option for trimming. The resulting dataset was subsequently analyzed for divergence time estimation using MCMCTREE, implemented in PAML version 4.10.6 (Yang, 2007). To ensure convergence, two independent MCMCTree runs were conducted, each with a burn-in of 100,000 iterations. Markov chains were sampled every 1,000 iterations until a total of 20,000 trees were produced, employing the approximate likelihood algorithm. Prior distributions for sigma² and rgene were set to G(1, 10) and G(1, 50), respectively, based on substitution rates inferred from BaseML. Reference-calibrated time points established using the TimeTree database (http://timetree.org/) included a fixed maximum age of 297.9 million years ago (Mya) for the Teleost infraclass. Additional nodes - Clupeocephala, Acanthomorphata, Euacanthomorphaceae, and Scombridae - were assigned minimum ages of 180, 128, 99.5, and 32.6 Mya, along with maximum ages of 251.5, 165, 116.7, and 58.3 Mya, respectively.

### Identification of Rapidly Evolving Gene Families

Rapidly evolving gene families were identified using CAFÉ version 5.1.1, which employs a stochastic birth and death model to characterize gene family evolution over time and detect significant size changes (Mendes et al., 2020). Viterbi p-values, calculated with a default significance threshold of 0.01, were utilized to assess significant contractions or expansions of gene families across branches. The birth-death parameter (lambda) was estimated using both single and multiple lambdas, with varying numbers of gamma rate categories (1-3) to ensure convergence; three independent runs were executed for each estimate. Parameter selection was guided by the best model final likelihood. Additionally, a customized R script was developed to extract and analyze gene families undergoing rapid evolution in extant species, utilizing R version 4.2

### *Detecting* Positive Selection of Genes

Positive selection was examined using CodeML from the PAML package (Yang, 2007), assessing all one-to-one orthologous genes under branch models with heterogeneous ω across branches. The northern bluefin tuna branch, including both T. thynnus and T. orientalis, was designated as the foreground branch for testing. Significance was evaluated through a likelihood ratio test.

### Trait Association Analysis

To identify genetic markers associated with trait variation within the polyphyletic bluefin tuna group, PhyloGWAS (Pease et al., 2016) was employed across the Scombridae lineage. PhyloGWAS assesses whether an excess of nonsynonymous variants is shared among individuals that are not monophyletic but exhibit a shared phenotypic trait. The analysis commenced with the download of raw FASTQ files for each Scombridae accession, followed by preprocessing with fastp (Chen, 2023; Chen et al., 2018) and HiFiAdapterFilt (Sim et al., 2022). Each sample was aligned to the main 24 chromosomes of the assembled T. thynnus genome using minimap2 (Li, 2021). Sequence variations were called using BCFtools (Danecek et al., 2021), and MVFtools (Pease & Rosenzweig, 2018) were employed to identify nonsynonymous variants correlated with bluefin tunas. The significance of these findings was evaluated by comparing the number of nonsynonymous variants with the expected count resulting from incomplete lineage sorting (ILS) alone, simulating a single change across 109 loci using ms over the consensus phylogeny.

## Results

The resultant phased genome assembly achieved contig/scaffold N50 values of 6.66/33.67 Mbp and 6.89/33.09 Mbp for the two haplotype assemblies, with ungapped sequence lengths of 859.8 Mbp and 858.6 Mbp, consisting mainly of 24 chromosomal scaffolds that encompass 97.5% of complete BUSCO genes for both haplotypes.

### HiFi and Omni-C reads enabled Phased assembly of Atlantic Bluefin Tuna genome

For the Atlantic bluefin tuna, a total of 2,391,784 CCS reads were generated with a mean read length of 22,042 bp. This yielded approximately 52.7Gb of sequence data used in the assembly. An initial assembly of ABFT HiFi reads resulted in contig N50 of 12.0 Mbp with total assembly size of 799.4 Mbp. Since this assembly had haplotype sequences collapsed, it had alleles representing only one of the two haplotypes. These contigs were then polished with HiFi reads and scaffolded with Omni-C reads using HiRise scaffolding algorithm resulting in a total sequence of 799.4 Mbp and an N50 of 34.33 Mbp. Next, phased HiFi reads were obtained by running a haplotype assembly pipeline on the scaffolds. Because there is no parental information available for the sequenced sample, phasing accuracy could not be determined by direct comparison. Instead, haplotypes separation was validated based on k-mer abundance in each haplotype. K-mers were found at > 1X frequency in the collapsed genome, whereas duplicated k-mers were depleted in the phased haplotypes. Heterozygous markers are also separated across the two haplotypes. The phased HiFi reads were assembled into each haplotype resulting in a contig N50 of 6.6 Mbp and 6.8 Mbp with total assembly size of 859.80 Mbp and 858.5 Mbp respectively. When scaffolding each haplotype with haplotype specific Omni-C reads, the resulting scaffold N50 was 33.7 Mbp and 33.1 Mbp, respectively, indicating both the haplotypes are scaffolded to chromosome-scale sequences (Table 1).

**Table 1:** Genomic Assembly Metrics for Tbuoous Species.

|  | *Thunnus thynnus*<br>Haplotype 1 | *Thunnus thynnus*<br>Haplotype 2 | *Thunnus orientalis* | *Thunnus maccoyii* | *Thunnus albacares* | *Thunnus orientalis* |
| --- | --- | --- | --- | --- | --- | --- |
| NCBI SRA Reference Number | This publication | This publication | This publication | GCF_910596095.1 | GCA_914725855.1 | GCA_021601225.1 |
| Number of Scaffolds | 266 | 135 | 474 | 58 | 69 | 948 |
| Total Scaffolded Assembly Size (bp) | 805,000,461 | 796,394,621 | 809,715,861 | 782,423,979 | 792,117,861 | 826,834,687 |
| Largest Scaffold Size (bp) | 42,079,636 | 41,950,415 | 41,359,765 | 41,002,747 | 41,207,429 | 34,580,358 |
| Scaffold N<sub>10</sub> Size (bp) | 34,107,232 | 34,196,657 | 34,062,839 | 33,761,101 | 34,622,995 | 13,192,983 |
| Scaffold N<sub>50</sub> Size (bp) | 26,827,212 | 26,893,001 | 21,305,290 | 26,533,419 | 26,463,036 | 244,769 |
| Number of Contigs | 359 | 216 | 490 | 153 | 139 | 1,337 |
| Sum of Contig Lengths (bp) | 804,991,161 | 796,386,521 | 809,714,261 | 782,392,479 | 792,094,561 | 826,795,787 |
| Largest Contig Size (bp) | 35,337,377 | 41,361,610 | 41,359,765 | 34,584,767 | 37,025,965 | 18,078,587 |
| Contig N50 Size (bp) | 20,273,516 | 21,213,696 | 30,786,244 | 26,803,536 | 26,795,544 | 4,079,918 |
| Contig N95 Size (bp) | 1,550,044 | 1,840,282 | 6,040,732 | 2,313,259 | 3,368,304 | 129,327 |
| Number of Gaps | 93 | 81 | 16 | 95 | 70 | 389 |
| Total Size of Gaps (bp) | 9,300 | 8,100 | 1,600 | 31,500 | 23,300 | 38,900 |

### Contiguity and Completeness of HiFi based tuna assemblies

In general, while the contig statistics are similar for all the assemblies shown, the assemblies generated with PacBio HiFi reads in combination with Dovetail Hi-C/Omni-C have larger scaffolds. The Scaffold N50 are all greater than 30Mb (Table 1) being 5 times longer to the scaffolds assembled generated with lower coverage continuous long reads (Suda et al., 2019). For the ABFT assemblies > 99.99% of the sequence is represented by an ungapped sequence further demonstrating the assembly power of these data types. Interestingly, a large portion (>90.0%) of ABFT sequences across the different assemblies is contained in 24 largest scaffolds which is the predicted number of tuna chromosomes from previous genetic map linkage studies on *T. orientalis*. (Uchino et al., 2018) The Hi-C contact map for scaffolds doesn’t indicate any signal for over-joining or fragmented sequence implying the resulting scaffolds are chromosome-scale.

### New tuna assemblies are more complete with respect to gene content

The two phased and collapsed ABFT assemblies harbor 97.5 and 97.6% of complete BUSCO genes, respectively (Table 2). The fragmented genes correspond to 1.2, 1.11, and 1.0% and the missing genes to 1.26, 1.35, and 1.3%, respectively (Table 2). The collapsed assemblies using HiFi reads contained between 96.8% and 97.6% complete BUSCO genes as well as opposed to the published *T. orientalis* assembly (Suda et al., 2019) which only had 88.2% BUSCO genes with elevated fragmented and missing counts of 4.9% and 6.9%, respectively (Table 2). This validation of genome completeness was performed using a large set of orthologous 4,584 Actinopterygii genes (Hughes et al., 2018)

**Table 2:** BUSCOS Results using Actinopterygll Gene Set.

|  | *Thunnus thynnus*<br>Haplotype 1 | *Thunnus thynnus*<br>Haplotype 2 | *Thunnus thynnus*<br>Diploid<br>Haplotype 1 +<br>Haplotype 2 | *Thunnus*<br>*orientalis* | *Thunnus*<br>*maccoyii* | *Thunnus*<br>*albacares* | *Thunnus*<br>*orientalis* |
| --- | --- | --- | --- | --- | --- | --- | --- |
| NCBI SRA<br>Reference<br>Number | This publication | This publication | This publication | This publication | GCF_910596095.<br>1 | GCA_914725855.<br>1 | GCA_021601225.<br>1 |
| Complete<br>BUSCO Genes | 98.9% | 99.1% | 99.3% | 99.1% | 99.1% | 99.2% | 98.2% |
| Single Copy<br>BUSCO Genes | 98.2% | 98.5% | 1.4% | 98.7% | 98.5% | 98.7% | 95.7% |
| Duplicated<br>BUSCO Genes | 0.7% | 0.6% | 97.9% | 0.4% | 0.6% | 0.5% | 2.5% |
| Fragmented<br>BUSCO Genes | 0.4% | 0.4% | 0.3% | 0.3% | 0.3% | 0.2% | 0.8% |
| Missing BUSCO<br>Genes | 0.7% | 0.6% | 0.4% | 0.6% | 0.6% | 0.6% | 1.0% |

We were also able to assess gene content using a publicly available *T. orientalis* coding sequence (CDS) data set which was made publicly available in 2013 (Nakamura et al., 2013). The various HiFi read based assemblies contained between 99.2% and 99.9% of the total 26,433 *T.orientalis* CDSs (Table 2). The majority (∼99%) of the mapped sequences were presented in single copy (Table 2) which was not surprising since these represented 1N haploid sized assemblies. The previously published *T. orientalis* reference (Suda et al., 2019) displayed a lower representation of CDS sequences at 97.7%. As with the HiFi based assemblies, the majority of the CDS (97%) were represented in single copy since the assembly was also a haplotype collapsed version.

### Synteny

The analysis revealed very few sequence rearrangements in the chromosomes between the two Atlantic Bluefin tuna haplotypes and between the genomes of T. *thunnus* and T. *orientalis* (Figure 1). This finding is consistent with the stable karyotype and comparable diploid chromosome number of 48, characterized by a high proportion of telocentric chromosomes among extant Thunnus species (Lee et al., 2018).

**Figure 1:**
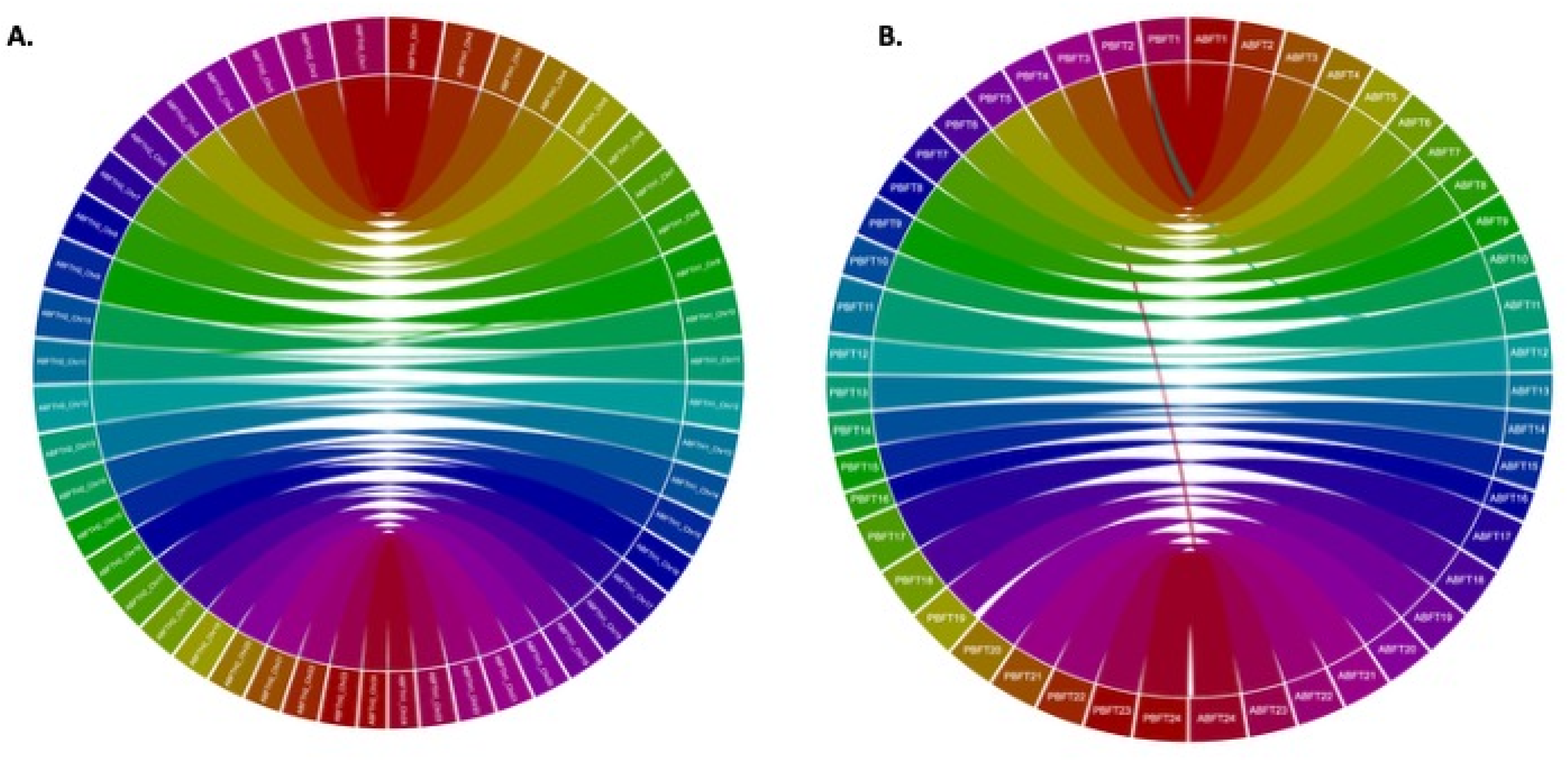
Synteny Analyses of ABFT phase 1 vs phase 2 (A) and ABFT vs PBFT (B) Synteny plot of alignments between ABFT phased chromosomal assemblies (A) and between ABFT haplotype 1 and PBFT chromosomes (B). Colors represent different chromosomes indicated by the chromosome names.

### Sequence comparison of phased haplotypes of the Atlantic bluefin tuna genome

The alignment of the two haplotypes revealed a sequence identity of 98.47%, which corresponds to an expected heterozygosity of 0.01 (approximately one difference every 100 bases) between the haplotypes. A more detailed comparison identified 2,369,313 single nucleotide variants (SNVs), 1,489 relocations, and 338 inversions. When comparing haplotype 2 against haplotype 1 as a reference, substantial numbers of biallelic variants were observed, including 502 kb of insertions, 495.3 kb of deletions, and 4.879 Mb of SNPs. In the analysis of multi-allelic variants, 1.268 kb of multi-allelic insertions, 1.722 kb of deletions, 68 instances of multi-allelic SNPs, and 1.1 kb of complex structural variations were identified. Reference calls accounted for 1.077 Mb of the haplotype 2 genome, and although they are included in the VCF file, their significance cannot be overlooked (Figure 2). The variant allele frequency by genotype demonstrated a predominance of heterozygous calls (0/x), with the most frequently observed changes being A to G, G to A, T to C, and C to T. Transition mutations were observed at a higher frequency (3.0 million) compared to transversions (∼1.9 million), indicating a bias towards transitions in the genetic changes occurring between the haplotypes. The analysis of small insertions and deletions showed a notable prevalence of insertions up to 400 bp, suggesting a dynamic structural variation landscape within these genomes (Supplementary Figure 1). Mapping depths for readings typically exceeded 50X coverage, displaying a Gaussian distribution characterized by a high mapping quality (Q60), which resulted in an overall high average genotype quality across the analyzed variants.

**Figure 2:**
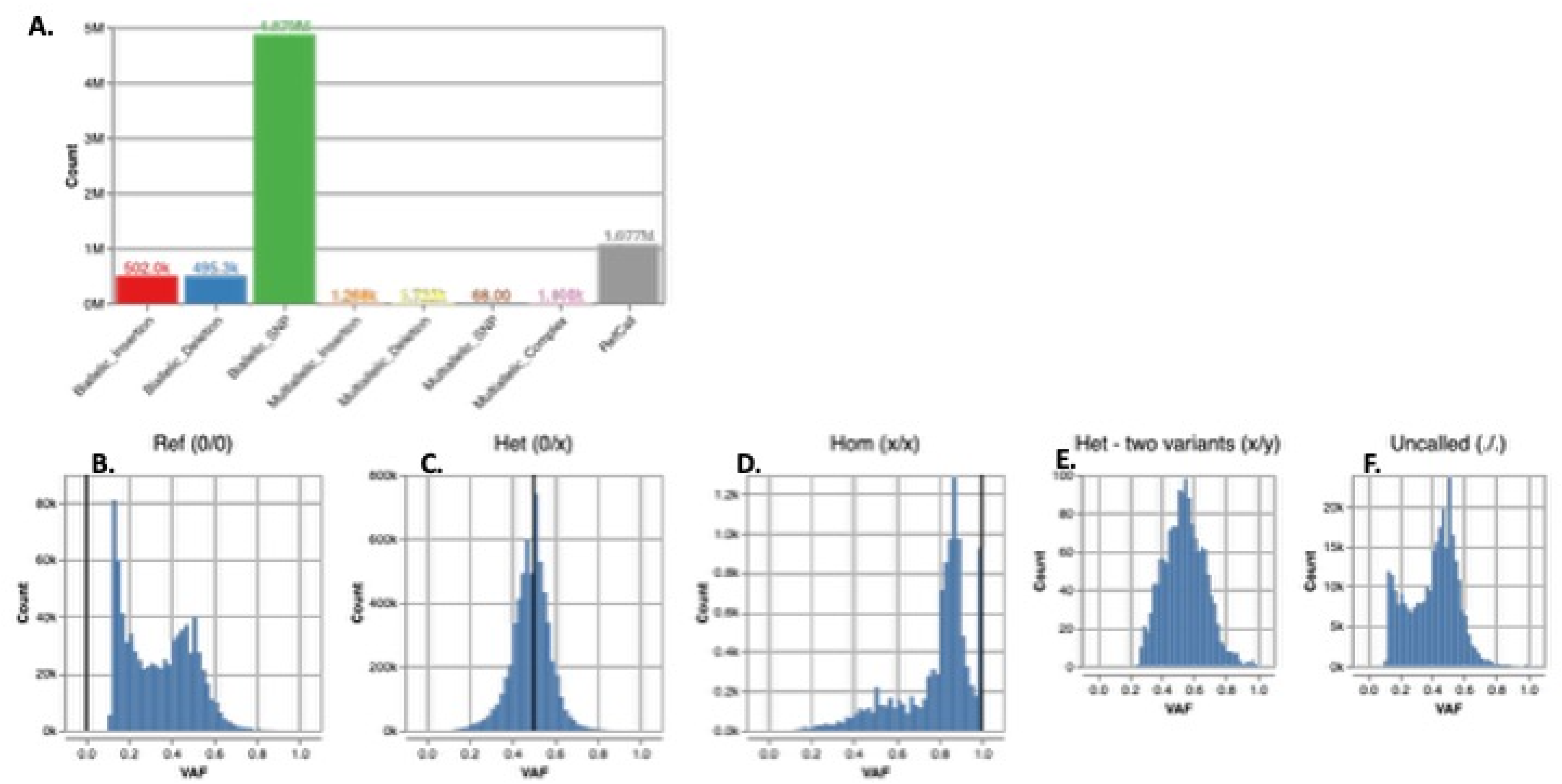
ABFT Hap 1 & Hap 2 variant comparison. **A)**. Percentage and type of biallelic and multi-allelic variants in ABFT haplotype 2, with haplotype 1 as reference, **B)** Variant allele frequency (VAF) of Reference (0/0), **C)** of all heterozygous to reference genotypes (0/x) **D)**., of all homozygous genotypes ( x/x) **E)**., all heterozygous-doubly variant genotypes (x/y) **F)**., and of all uncalled genotypes (-/-).

### Sequence comparison of Atlantic Bluefin Tuna with Pacific Bluefin Tuna genome

The comparison of the Atlantic Bluefin Tuna assembly with the recently published improved genome assembly of Pacific Bluefin Tuna (Suda et al., 2019) indicates that the assembly of Atlantic Bluefin Tuna, generated using HiFi reads, possesses a more comprehensive gene content compared to the latest Pacific Bluefin Tuna assembly. When haplotype 2 was compared to haplotype 1 as a reference, the analysis identified 727 kb of biallelic insertions, 756 kb of deletions, and 7.937 Mb of SNPs (Table 3). Among multi-allelic variants, 50.69 kb, 30.65 kb, and 57.13 kb of multi-allelic insertions, deletions, and SNPs, as well as complex variants, were recorded. Reference calls accounted for 1.646 Mb of the Pacific Bluefin Tuna genome (Suda et al., 2019). The variant allele frequency distribution revealed a majority of heterozygous calls (0/x), with common changes occurring from A to G, G to A, T to C, and C to T. A higher frequency of transitions (4.898 million) compared to transversions (3.039 million) was observed, resulting in a Transition/Transversion ratio of 1.61, which is similar to the ratio (1.63) found when comparing the two Atlantic Bluefin Tuna haplotypes. Additionally, a notable prevalence of insertions up to 400 bp was reported in the analysis of small insertions and deletions.

**Table 3:** Gene Mapping Results using *Thunnus maccoyil* Annotation Gene Set.

|  | Thunnus thynnus<br>Haplotype 1 | Thunnus thynnus<br>Haplotype 2 | Thunnus<br>orientalis | Thunnus maccoyii | Thunnus<br>orientalis |
| --- | --- | --- | --- | --- | --- |
| NCBI SRA<br>Reference<br>Number | This publication | This publication | This publication | GCF_910596095.1 | GCA_021601225.1 |
| Thunnus maccoyii<br>Transcript<br>Complete Full<br>Length | 95.9% | 95.9% | 96.0% | 97.7% | 97.0% |
| Thunnus maccoyii<br>Transcript<br>Fragmented | 2.2% | 2.2% | 2.1% | 0.7% | 3.0% |
| Thunnus maccoyii<br>Transcript Missing | 1.9% | 1.9% | 1.9% | 1.6% | >>0.1% |

### Genome annotation

The repeat elements of T. thynnus exhibit a GC content of approximately 39.83%, which is comparable to the genome-wide GC content of 40%. Repetitive regions comprise 27.99% (225,337,774 bp) of the total genome length. Among these repetitive elements, DNA repetitive elements are the most prevalent, accounting for 11.44%, followed by LINEs at 4.4% and LTRs at 1.4%. Small RNAs, satellites, and simple repeats show lower representation, with 0.37%, 0.21%, and 0.18%, respectively. The annotation results indicate a total of 27,859 genes and 35,273 transcripts. More than 98% of the gene sequences displayed at least one hit in the RefSeq or SwissProt databases, while 93% of the gene sequences matched with one or more databases in the InterProScan analysis. Importantly, approximately 18,528 protein sequences were mapped to a SUPERFAMILY of proteins, 22,373 had at least one protein domain, and 1,318 proteins are involved in established KEGG pathways (Table 4).

**Table 4:**
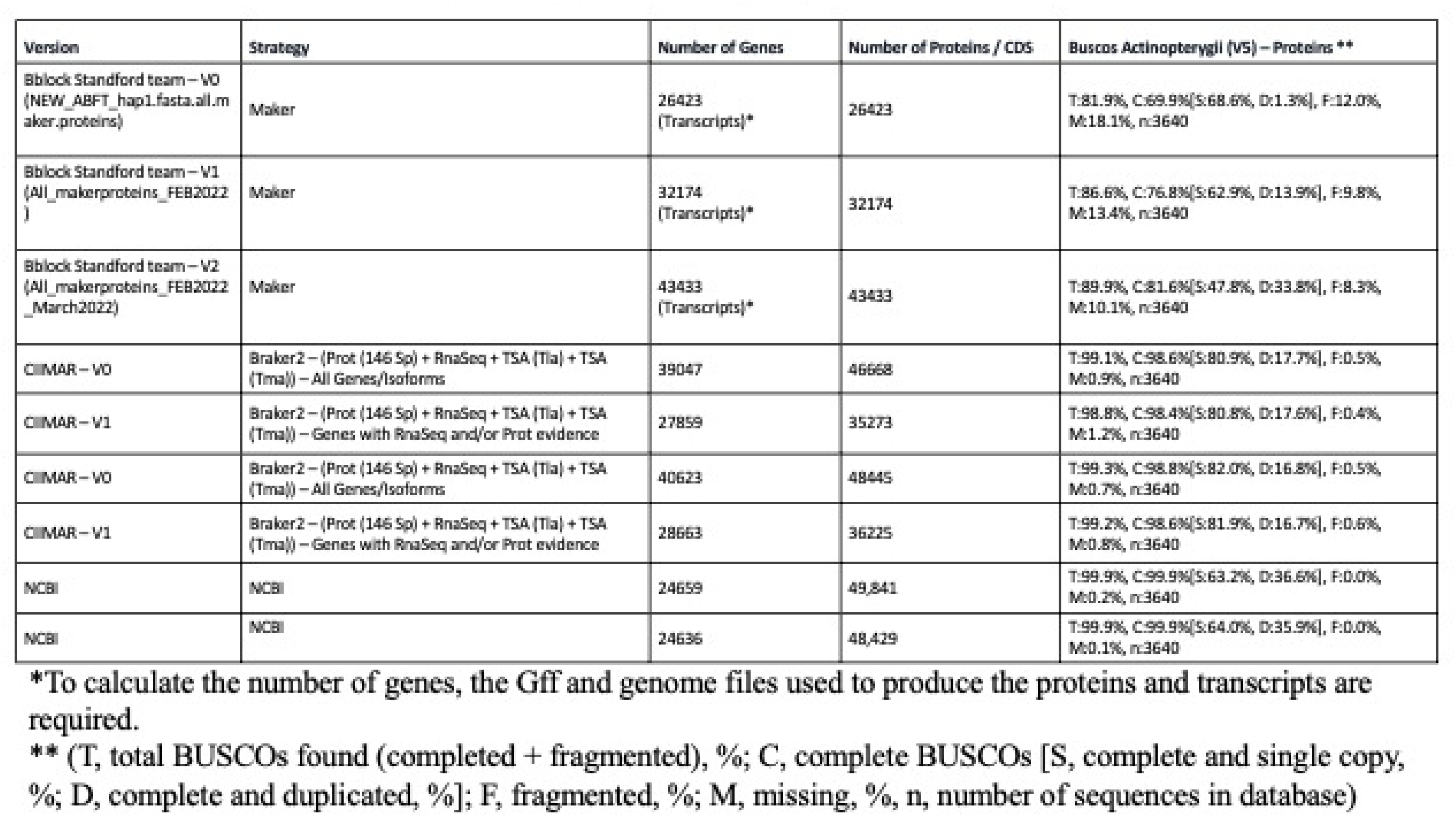
ABFf and PBFf annotation versus other tuna species.

### Topology Weighting analysis

The most frequent topology across the genome was the ‘species-tree’ topology, with an average weighting of 0.37, followed by the ‘bluefin-tree’ topology, which had an average weighting of 0.29. The only other topology with a genome-wide weighting greater than 0.05 was one in which the Yellowfin tuna was more closely related to northern bluefin tunas than to southern bluefin tunas, with a weighting of 0.19. Generally, the ‘species-tree’ topology was predominant at the beginning and end of linkage groups, while the ‘bluefin-tree’ topology appeared slightly more frequently in the middle of the linkage groups (Figure 3). The results were similar regardless of whether the Atlantic Bluefin Tuna (ABFT) or Pacific Bluefin Tuna (PBFT) was used as the reference.

**Figure 3:**
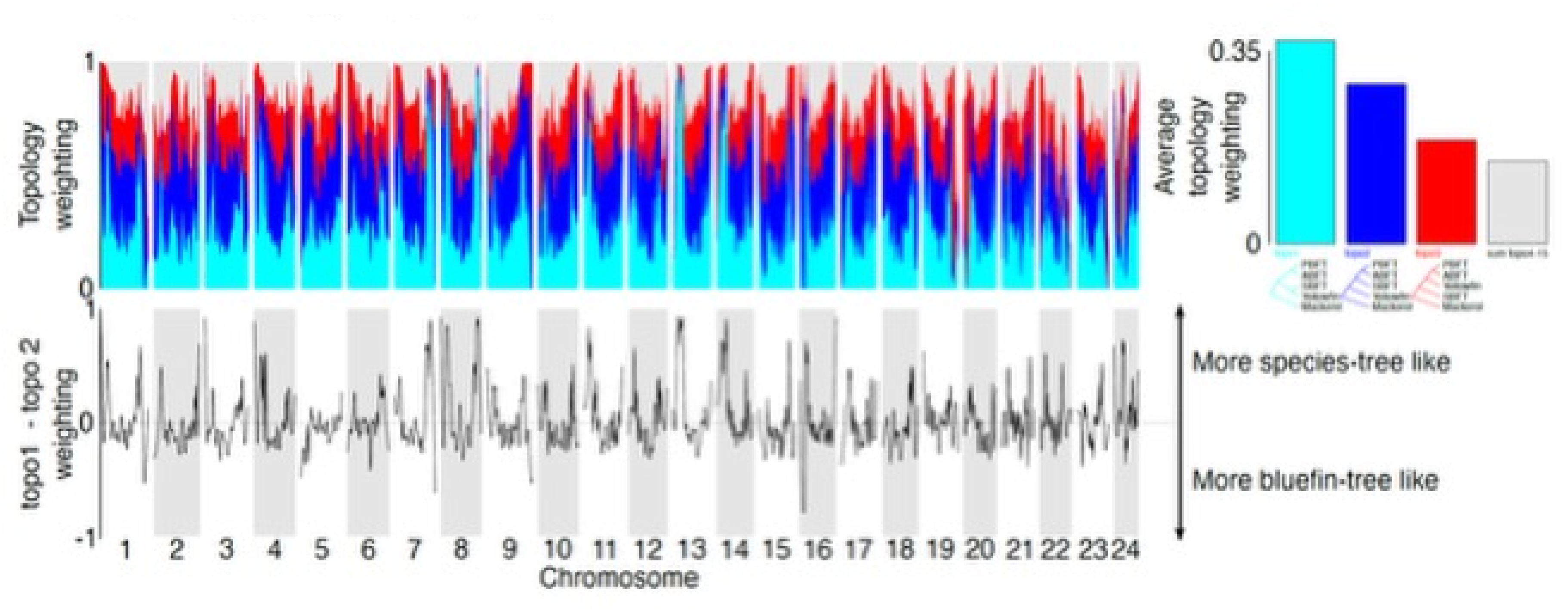
Topology weighting analysis. Topology weighting analysis (TWISST) across the 24 ABFT genome assembly chromosomes. Turquoise colours represent the species tree; blue the “bluefin”, and red the only other frequent topology, the remaining 12 topologies are coloured in grey. Bottom row shows the weighting of the bluefin tree subtracted from the weighting of the species-tree; regions where the line is positive (above the dotted line), generally support the species-tree, whereas below the line support the bluefin tree.

**Figure 4:**
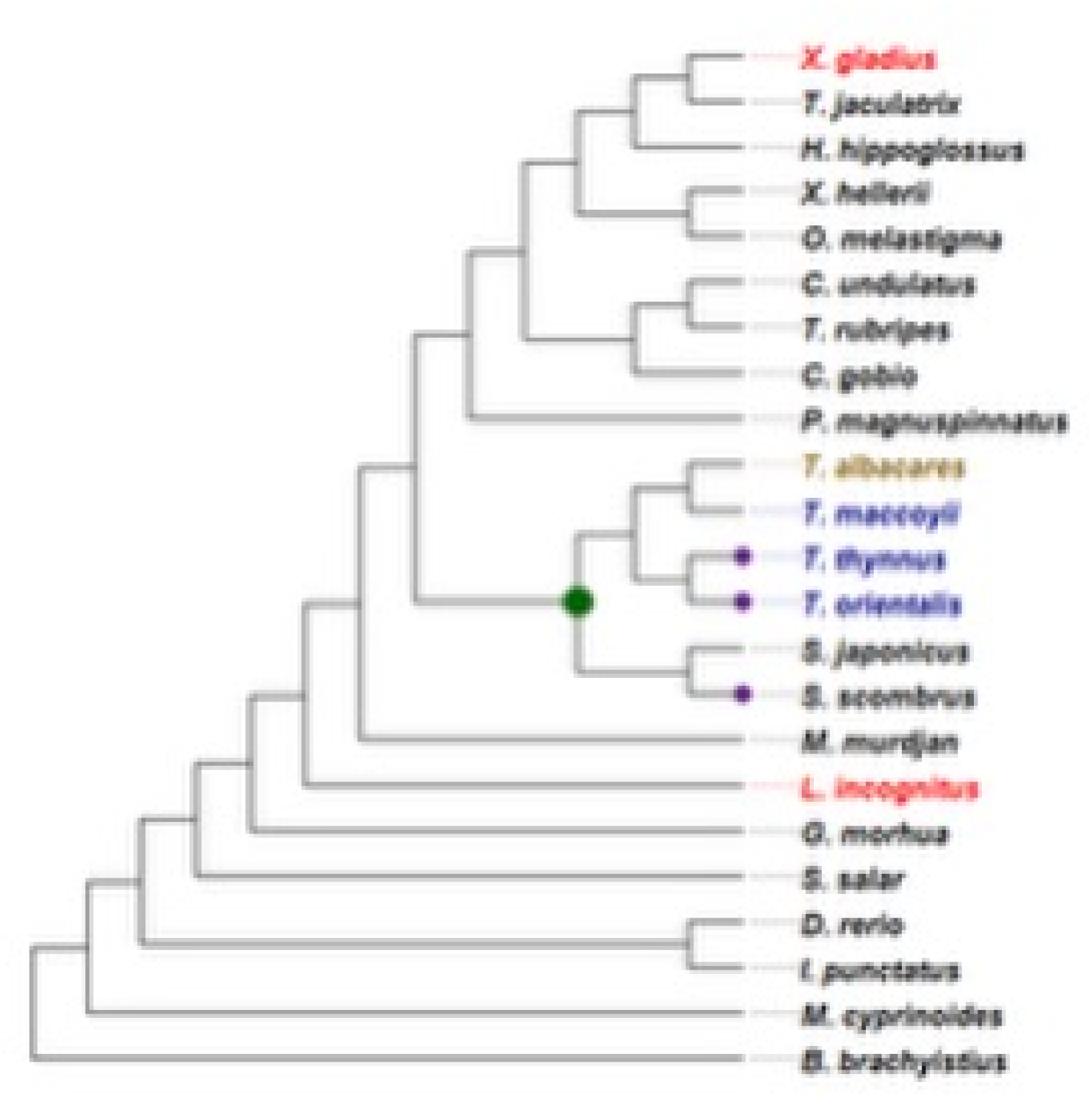
Phylogenetic analyses of Scombridae using orthologous genes. Phylogenetic relationships among scombrid and related teleost species based on mitochondrial genome sequences. The tree illustrates the placement of **T. thynnus**, **T. orientalis**, and **T. maccoyii** (highlighted in purple) within the *Thunnus*clade, showing their close evolutionary relationships. The green dot indicates the most recent common ancestor of the *Thunnus* and *Scomber* lineages. *T. albacares* (gold) serves as an outgroup to the bluefin tuna species. Other teleost representatives are shown for context, including *Xiphias gladius* and *Lutjanus incognitus* (in red) as more distantly related taxa. Branch colors correspond to major taxonomic groupings used in comparative analyses.

The genome of T. thynnus comprises 27.99% mobile elements. This proportion of repeat elements aligns with findings from other scombriform species. In genome projects for *Scomber colias* and *Euthynnus affinis*, members of the Scombridae family, the reported percentages of repetitive elements were 29.62% and 25.59%, respectively (Lein et al., 2007; Machado et al., 2021). Additionally, the genome annotation of closely related species, *Thunnus maccoyii* and *Thunnus albacares*, revealed 26.07% and 26.27% of repetitive sequences (“*Thunnus albacares* Annotation Report,” 2021; “Thunnus maccoyii Annotation Report,” 2021). In the ABFT genome, DNA transposons represented the most significant proportion of the repeat content, while LTRs and SINEs were the least represented. These patterns have been consistently observed in other teleost species (Gao et al., 2016; Shao et al., 2019).

A high level of polymorphism was observed, with a total of 4,879,068 biallelic SNPs across the two haplotypes of the phased T. thynnus genome. Consequently, the two haplotypes exhibit approximately 1% heterozygosity and 98.5% homozygosity, confirming the highly polymorphic nature of T. thynnus, which is consistent with a large population size and a wide geographical distribution across the Atlantic. This level of heterozygosity is significantly higher compared to vertebrate populations that have experienced a bottleneck (E. E. Armstrong et al., 2020). These findings suggest that the T. thynnus population may possess sufficient variation and elevated heterozygosity to confer a fitness advantage, assuming that polymorphisms contribute to improved fitness, and may serve as a valuable resource for adaptation to natural selection. Nevertheless, several distinct populations are thought to exist within the Atlantic basin and adjacent seas, highlighting the need for population-level assessments of heterozygosity to accurately evaluate the true levels of genetic diversity within each known breeding population.

### Orthology Analysis

The orthology analysis focused on the comparison of *Thunnus* species within the Scombridae family and their evolutionary relationship with the outgroup Homo sapiens. Among the evaluated species, *Thunnus orientalis* and *Thunnus thynnus* exhibited gene counts of 28,636 and 27,843 genes, respectively, with 96.2% and 96.9% of these genes categorized within 20,167 and 19,978 orthogroups. In comparison, *Homo sapiens* displayed a total of 20,065 genes, with 92.6% in 14,230 orthogroups, indicating significant genetic divergence between humans and teleosts. Notably, Salmo salar contained 42,985 genes, which contributes to its comparative analysis against *Thunnus* species, highlighting its extensive genomic capacity . The results underscore the shared genetic heritage within the Scombridae family and illustrate how gene family expansions may contribute to the adaptive strategies of bluefin tunas in response to environmental pressures (Table 5).

**Table 5:**
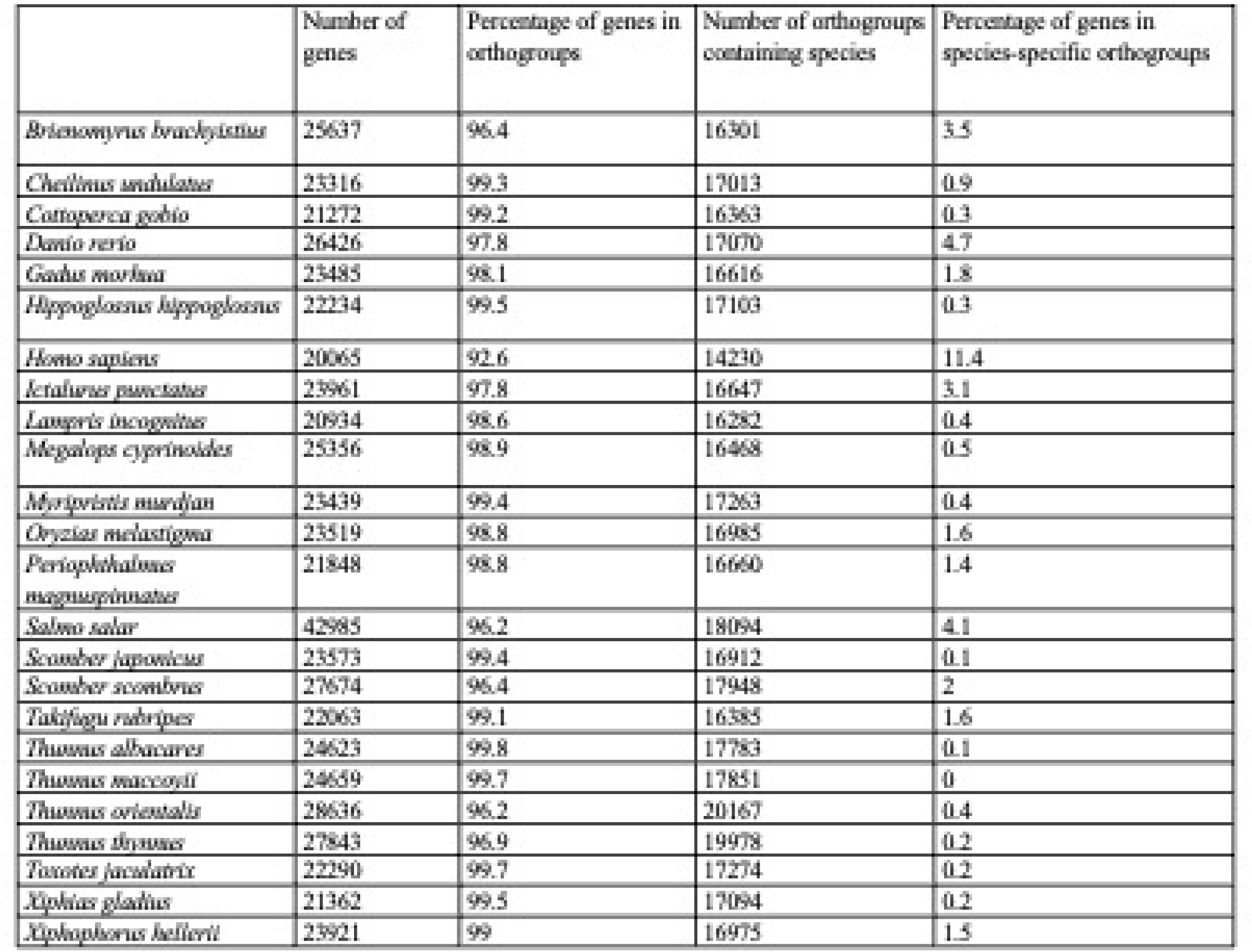
Orthology analysis.

### Results of Functional Annotation and Gene Analysis

The functional annotation of genes across the evaluated teleost species, particularly within the Scombridae family, revealed a rich diversity of biological functions. Notable genes such as IFI44L were identified as interferon-induced proteins, suggesting a potential role in mediating immune responses to environmental stressors. Similarly, PLXNC1 was associated with signal transduction, indicating its involvement in critical intracellular communication pathways.

Furthermore, the functional profiles demonstrated strong representation of various molecular activities, including adenylate cyclase activity and GABA-A receptor activity, which are vital for cellular regulation and response mechanisms. High levels of voltage-gated potassium channel activity were also detected, highlighting its significance in maintaining neuronal excitability and muscle contraction.

The data also indicated that five genes were specifically identified as significant between the analyses of gene variation and expansion. This limited number suggests a focused adaptive evolution within select gene families, emphasizing their potential importance in the adaptive radiation of bluefin tunas and their relatives.

## Conclusion

A complete chromosome-scale genome assembly for Atlantic bluefin tuna (*Thunnus thynnus*), a highly sought-after commercial species, has been achieved using Pacific Biosciences HiFi sequencing technology. This assembly leverages circular consensus sequencing, which employs multiple passes around a circularized template to generate highly accurate long-read consensus sequences. The precision of this sequencing technique significantly enhances the ability to construct an accurate genome assembly. This reference genome holds considerable promise for the detection of various types of genetic variants within Atlantic bluefin populations across the Atlantic and adjacent seas, thereby facilitating improved population-level management strategies. Furthermore, the comprehensive analysis highlights the functional relevance of numerous orthologs, shedding light on the evolutionary dynamics and ecological resilience of teleost species. By elucidating specific gene functions and their distributions, these findings deepen the understanding of the molecular mechanisms that drive the adaptations of bluefin tunas, ultimately informing conservation efforts and guiding future research in evolutionary biology.

Figure S1: **A)**. Coverage depth distribution of variants, **B)**. Quality distribution of variants, **C)**. Range of genotype quality across all genotypes detected, **D)**. Count of single nucleotide polymorphism for each reference nucleotide -A, G, T, C., **E)**. Count of transitions (Ti) and transversion (Tv) events with respect to the reference with a biallelic Ti/Tv ratio of 1.61. and , **F)**. Biallelic indel size distribution.

## References

1. Alexa, A., & Rahnenführer, J. (2009). Gene set enrichment analysis with topGO. http://www.mpi-sb.mpg.de/∼alexa

2. Armstrong, E., Taylor, R. W., Miller, D. E., Kaelin, C. B., Barsh, G. S., Hadly, E. A., & Petrov, D. (2020). Long live the king: Chromosome-level assembly of the lion (Panthera leo) using linked-read, Hi-C, and long-read data. BMC Biology, 18(1). 10.1186/s12915-019-0734-5

3. Armstrong, J., Hickey, G., Diekhans, M., Fiddes, I. T., Novak, A. M., Deran, A., Fang, Q., Xie, D., Feng, S., Stiller, J., Genereux, D., Johnson, J., Marinescu, V. D., Alföldi, J., Harris, R. S., Lindblad-Toh, K., Haussler, D., Karlsson, E., Jarvis, E. D., … Paten, B. (2020). Progressive Cactus is a multiple-genome aligner for the thousand-genome era. Nature, 587(7833), 246–251. 10.1038/s41586-020-2871-y

4. Arrizabalaga, H., Dufour, F., Kell, L., Merino, G., Ibaibarriaga, L., Chust, G., Irigoien, X., Santiago, J., Murua, H., Fraile, I., Chifflet, M., Goikoetxea, N., Sagarminaga, Y., Aumont, O., Bopp, L., Herrera, M., Fromentin, J., & Bonhomeau, S. (2015). Global habitat preferences of commercially valuable tuna. Deep-Sea- Research II, 113, 102–112. 10.1016/j.dsr2.2014.07.001

5. Bernos, T. A., Jeffries, K. M., & Mandrak, N. E. (2020). Linking genomics and fish conservation decision making: a review. In Reviews in Fish Biology and Fisheries (Vol. 30, Issue 4, pp. 587–604). Springer Science and Business Media Deutschland GmbH. 10.1007/s11160-020-09618-8

6. Blank, J. M., Farwell, C. J., Morrissette, J. M., Schallert, R. J., & Block, B. A. (2007). Influence of Swimming Speed on Metabolic Rates of Juvenile Pacific Bluefin Tuna and Yellowfin Tuna. In Physiological and Biochemical Zoology (Vol. 80, Issue 2).

7. Block, B. A., & Finnerty, J. R. (1994). Endothermy in fishes: a phylogenetic analysis of constraints, predispositions, and selection pressures. Environmental Biology of Fishes, 40(3), 283–302. 10.1007/BF00002518

8. Block, B. A., Whitlock, R., Schallert, R. J., Wilson, S., Stokesbury, M. J. W., Castleton, M., & Boustany, A. (2019). Estimating Natural Mortality of Atlantic Bluefin Tuna Using Acoustic Telemetry. Scientific Reports, 9(1). 10.1038/s41598-019-40065-z

9. Block, B., Dewar, H., Farwell, C., & Princes, E. (1998). A new satellite technology for tracking the movements of Atlantic bluefin tuna. Proceedings of the National Academy of Sciences, 95, 9384–9389.

10. Block, B., & Stevens, D. (2001). Tuna Physiology, Ecology and Evolution. In W. Hoar, D. Randall, & A. Farell (Eds.), Fish Physiology (Vol. 19). Academic Press.

11. Brůna, T., Hoff, K. J., Lomsadze, A., Stanke, M., & Borodovsky, M. (2021). BRAKER2: Automatic eukaryotic genome annotation with GeneMark-EP+ and AUGUSTUS supported by a protein database. NAR Genomics and Bioinformatics, 3(1), 1–11. 10.1093/nargab/lqaa108

12. Buchfink, B., Xie, C., & Huson, D. H. (2014). Fast and sensitive protein alignment using DIAMOND. In Nature Methods (Vol. 12, Issue 1, pp. 59–60). Nature Publishing Group. 10.1038/nmeth.3176

13. Cantalapiedra, C. P., Hern̗andez-Plaza, A., Letunic, I., Bork, P., & Huerta-Cepas, J. (2021). eggNOG-mapper v2: Functional Annotation, Orthology Assignments, and Domain Prediction at the Metagenomic Scale. Molecular Biology and Evolution, 38(12), 5825–5829. 10.1093/molbev/msab293

14. Cantarel, B. L., Korf, I., Robb, S. M. C., Parra, G., Ross, E., Moore, B., Holt, C., Alvarado, A. S., & Yandell, M. (2008). MAKER: An easy-to-use annotation pipeline designed for emerging model organism genomes. Genome Research, 18(1), 188–196. 10.1101/gr.6743907

15. Capella-Gutiérrez, S., Silla-Martínez, J. M., & Gabaldón, T. (2009). trimAl: A tool for automated alignment trimming in large-scale phylogenetic analyses. Bioinformatics, 25(15), 1972–1973. 10.1093/bioinformatics/btp348

16. Chen, S. (2023). Ultrafast one-pass FASTQ data preprocessing, quality control, and deduplication using fastp. IMeta, 2(2). 10.1002/imt2.107

17. Chen, S., Zhou, Y., Chen, Y., & Gu, J. (2018). Fastp: An ultra-fast all-in-one FASTQ preprocessor. Bioinformatics, 34(17), i884–i890. 10.1093/bioinformatics/bty560

18. Ciezarek, A. G., Osborne, O. G., Shipley, O. N., Brooks, E. J., Tracey, S. R., Mcallister, J. D., Gardner, L. D., Sternberg, M. J. E., Block, B., & Savolainen, V. (2019). Phylotranscriptomic Insights into the Diversification of Endothermic Thunnus Tunas. Molecular Biology and Evolution, 36(1), 84–96. 10.1093/molbev/msy198

19. Collette, B., Boustany, A., Fox, W., Graves, J., Juan Jorda, M., & Restrepo. V. (2021). Thunnus thynnus: Collette, B.B., Boustany, A., Fox, W., Graves, J., Juan Jorda, M. &amp; Restrepo, V. In IUCN Red List of Threatened Species. 10.2305/IUCN.UK.2021-2.RLTS.T21860A46913402.en

20. Dainat J. (2021). AGAT: Another Gff Analysis Toolkit to handle annotations in any GTF/GFF format.

21. Danecek, P., Bonfield, J. K., Liddle, J., Marshall, J., Ohan, V., Pollard, M. O., Whitwham, A., Keane, T., McCarthy, S. A., & Davies, R. M. (2021). Twelve years of SAMtools and BCFtools. GigaScience, 10(2). 10.1093/gigascience/giab008

22. Darling, A. E., Mau, B., & Perna, N. T. (2010). Progressivemauve: Multiple genome alignment with gene gain, loss and rearrangement. PLoS ONE, 5(6). 10.1371/journal.pone.0011147

23. Díaz-Arce, N., Arrizabalaga, H., Murua, H., Irigoien, X., & Rodríguez-Ezpeleta, N. (2016). RAD-seq derived genome-wide nuclear markers resolve the phylogeny of tunas. Molecular Phylogenetics and Evolution, 102, 202–207. 10.1016/j.ympev.2016.06.002

24. Dimens, P. V, Hildahl, T., Mcpeak, M. B., Jones, K. L., Cusatti, S., Margulies, D., Scholey, V., & Saillant, E. A. (2024). Genomic resources for the yellowfin tuna nus albacares. Molecular Biology Reports, 946. 10.1007/s11033-023-09117-6

25. Emms, D. M., & Kelly, S. (2019). OrthoFinder: Phylogenetic orthology inference for comparative genomics. Genome Biology, 20(1). 10.1186/s13059-019-1832-y

26. Fritsches, K. A., Brill, R. W., & Warrant, E. J. (2005). Warm Eyes Provide Superior Vision in Swordfishes ters of up to 90 mm (Figure 1A). Such large eyes suggest. Current Biology, 15, 55–58. 10.1016/j

27. Fujioka, K., Masujima, M., Boustany, A. M., & Kitagawa, T. (2015). Horizontal Movements of Pacific Bluefin Tuna.

28. Gao, B., Shen, D., Xue, S., Chen, C., Cui, H., & Song, C. (2016). The contribution of transposable elements to size variations between four teleost genomes. Mobile DNA, 7(1). 10.1186/s13100-016-0059-7

29. Garg, S., Fungtammasan, A., Carroll, A., Chou, M., Schmitt, A., Zhou, X., Mac, S., Peluso, P., Hatas, E., Ghurye, J., Maguire, J., Mahmoud, M., Cheng, H., Heller, D., Zook, J. M., Moemke, T., Marschall, T., Sedlazeck, F. J., Aach, J., … Li, H. (2021). Chromosome-scale, haplotype-resolved assembly of human genomes. Nature Biotechnology, 39(3), 309–312. 10.1038/s41587-020-0711-0

30. Hoff, K. J., Lange, S., Lomsadze, A., Borodovsky, M., & Stanke, M. (2016). BRAKER1: Unsupervised RNA-Seq-based genome annotation with GeneMark-ET and AUGUSTUS. Bioinformatics, 32(5), 767–769. 10.1093/bioinformatics/btv661

31. Hoff, K. J., Lomsadze, A., Borodovsky, M., & Stanke, M. (2019). Whole-genome annotation with BRAKER. In Methods in Molecular Biology (Vol. 1962, pp. 65–95). Humana Press Inc. 10.1007/978-1-4939-9173-0_5

32. Huerta-Cepas, J., Szklarczyk, D., Heller, D., Hernández-Plaza, A., Forslund, S. K., Cook, H., Mende, D. R., Letunic, I., Rattei, T., Jensen, L. J., Von Mering, C., & Bork, P. (2019). EggNOG 5.0: A hierarchical, functionally and phylogenetically annotated orthology resource based on 5090 organisms and 2502 viruses. Nucleic Acids Research, 47(D1), D309–D314. 10.1093/nar/gky1085

33. Hughes, L. C., Ortí, G., Huang, Y., Sun, Y., Baldwin, C. C., Thompson, A. W., Arcila, D., Betancur, R., Li, C., Becker, L., Bellora, N., Zhao, X., Li, X., Wang, M., Fang, C., Xie, B., Zhou, Z., Huang, H., Chen, S., … Performed, Q. S. (2018). Comprehensive phylogeny of ray-finned fishes (Actinopterygii) based on transcriptomic and genomic data. 115(24), 6249–6254. 10.5061/dryad

34. Jones, P., Binns, D., Chang, H. Y., Fraser, M., Li, W., McAnulla, C., McWilliam, H., Maslen, J., Mitchell, A., Nuka, G., Pesseat, S., Quinn, A. F., Sangrador-Vegas, A., Scheremetjew, M., Yong, S. Y., Lopez, R., & Hunter, S. (2014). InterProScan 5: Genome-scale protein function classification. Bioinformatics, 30(9), 1236–1240. 10.1093/bioinformatics/btu031

35. Juan-Jordá, M. J., Mosqueira, I., Cooper, A. B., Freire, J., & Dulvy, N. K. (2011). Global population trajectories of tunas and their relatives. Proceedings of the National Academy of Sciences of the United States of America, 108(51), 20650–20655. 10.1073/pnas.1107743108

36. Flynn, M., Robert Hubley, Clément Goubert, Jeb Rosen, Andrew G. Clark, Cédric Feschotte, & Arian F. Smit. (2020). RepeatModeler2 for automated genomic discovery of transposable element families. PNAS, 19(117). 10.1186/s13059-018-1577-z

37. Kiełbasa, S. M., Wan, R., Sato, K., Horton, P., & Frith, M. C. (2011). Adaptive seeds tame genomic sequence comparison. Genome Research, 21(3), 487–493. 10.1101/gr.113985.110

38. Kim, D., Langmead, B., & Salzberg, S. L. (2015). HISAT: A fast spliced aligner with low memory requirements. Nature Methods, 12(4), 357–360. 10.1038/nmeth.3317

39. Kim, H., Lee, K., Lim, D. Il, Nam, S. Il, Han, S. hee, Kim, J., Lee, E., Han, I. S., Jin, Y. K., & Zhang, Y. (2019). Increase in anthropogenic mercury in marginal sea sediments of the Northwest Pacific Ocean. Science of the Total Environment, 654, 801–810. 10.1016/j.scitotenv.2018.11.076

40. Klinger, D. H., & Mendoza, N. (2019). The resource and environmental intensity of bluefin tuna aquaculture. In B. Block (Ed.), The Future of Bluefin Tunas: Ecology, Fisheries Management, and Conservation (pp. 312–334).

41. Lee, Y. H., Yen, T. B., Chen, C. F., & Tseng, M. C. (2018). Variation in the karyotype, cytochrome b gene, and 5s rDNA of four thunnus (Perciformes, Scombridae) tunas. Zoological Studies, 57. 10.6620/ZS.2018.57-34

42. Lein, E. S., Hawrylycz, M. J., Ao, N., Ayres, M., Bensinger, A., Bernard, A., Boe, A. F., Boguski, M. S., Brockway, K. S., Byrnes, E. J., Chen, L., Chen, L., Chen, T. M., Chin, M. C., Chong, J., Crook, B. E., Czaplinska, A., Dang, C. N., Datta, S., … Jones, A. R. (2007). Genome-wide atlas of gene expression in the adult mouse brain. Nature, 445(7124), 168–176. 10.1038/nature05453

43. Li, H. (2021). New strategies to improve minimap2 alignment accuracy. Bioinformatics, 37(23), 4572–4574. 10.1093/bioinformatics/btab705

44. Machado, A. M., Gomes-dos-Santos, A., Fonseca, M., da Fonseca, R. R., Veríssimo, A., Felício, M., Capela, R., Alves, N., Santos, M., Salvador-Caramelo, F., Domingues, M., Ruivo, R., Froufe, E., & Castro, L. F. C. (2021). A genome assembly of the Atlantic chub mackerel (Scomber colias): a valuable teleost fishing resource. 10.1101/2021.11.19.468211

45. MacKenzie, K. M., Robertson, D. R., Adams, J. N., Altieri, A. H., & Turner, B. L. (2019). Structure and nutrient transfer in a tropical pelagic upwelling food web: From isoscapes to the whole ecosystem. Progress in Oceanography, 178(February), 102145. 10.1016/j.pocean.2019.102145

46. Maddison, W. P. (1997). Gene Trees in Species Trees. In Systematic Biology (Vol. 46, Issue 3).

47. Mapleson, D., Accinelli, G. G., Kettleborough, G., Wright, J., & Clavijo, B. J. (2017). KAT: A K-mer analysis toolkit to quality control NGS datasets and genome assemblies. Bioinformatics, 33(4), 574–576. 10.1093/bioinformatics/btw663

48. Mayakonda, A., Lin, D. C., Assenov, Y., Plass, C., & Koeffler, H. P. (2018). Maftools: Efficient and comprehensive analysis of somatic variants in cancer. Genome Research, 28(11), 1747–1756. 10.1101/gr.239244.118

49. McDowall, J., & Hunter, S. (2011). InterPro Protein Classification. In Bioinformatics for Comparative Proteomics (pp. 37–47). Humana Press. 10.1007/978-1-60761-977-2_3

50. Mendes, F. K., Vanderpool, D., Fulton, B., & Hahn, M. W. (2020). CAFE 5 models variation in evolutionary rates among gene families. Bioinformatics, 36(22–23), 5516–5518. 10.1093/bioinformatics/btaa1022

51. Nakamura, Y., Mori, K., Saitoh, K., Oshima, K., Mekuchi, M., Sugaya, T., Shigenobu, Y., Ojima, N., Muta, S., Fujiwara, A., Yasuike, M., Oohara, I., Hirakawa, H., Chowdhury, V. S., Kobayashi, T., Nakajima, K., Sano, M., Wada, T., Tashiro, K., … Inouye, K. (2013). Evolutionary changes of multiple visual pigment genes in the complete genome of Pacific bluefin tuna. Proceedings of the National Academy of Sciences of the United States of America, 110(27), 11061–11066. 10.1073/pnas.1302051110

52. Nakatsuka, S. (2020). Stock Structure of Pacific Bluefin Tuna (Thunnus orientalis) for Management Purposes—A Review of Available Information. In Reviews in Fisheries Science and Aquaculture (Vol. 28, Issue 2, pp. 170–181). Taylor and Francis Inc. 10.1080/23308249.2019.1686455

53. Nurk, S., Walenz, B. P., Rhie, A., Vollger, M. R., Logsdon, G. A., Grothe, R., Miga, K. H., Eichler, E. E., Phillippy, A. M., & Koren, S. (2020). HiCanu: Accurate assembly of segmental duplications, satellites, and allelic variants from high-fidelity long reads. Genome Research, 30(9), 1291–1305. 10.1101/GR.263566.120

54. O’Leary, N. A., Wright, M. W., Brister, J. R., Ciufo, S., Haddad, D., McVeigh, R., Rajput, B., Robbertse, B., Smith-White, B., Ako-Adjei, D., Astashyn, A., Badretdin, A., Bao, Y., Blinkova, O., Brover, V., Chetvernin, V., Choi, J., Cox, E., Ermolaeva, O., … Pruitt, K. D. (2016). Reference sequence (RefSeq) database at NCBI: Current status, taxonomic expansion, and functional annotation. Nucleic Acids Research, 44(D1), D733–D745. 10.1093/nar/gkv1189

55. Ostrovsky, A., Hillman-Jackson, J., Bouvier, D., Clements, D., Afgan, E., Blankenberg, D., Schatz, M. C., Nekrutenko, A., Taylor, J., & Lariviere, D. (2021). Using Galaxy to Perform Large-Scale Interactive Data Analyses—An Update. Current Protocols, 1(2). 10.1002/cpz1.31

56. Pagniello, C. M. L. S., Maoiléidigh, N., Maxwell, H., Castleton, M. R., Aalto, E. A., Dale, J. J., Schallert, R. J., Stokesbury, M. J. W., Cosgrove, R., Dedman, S., Drumm, A., O’Neill, R., & Block, B. A. (2023). Tagging of Atlantic bluefin tuna off Ireland reveals use of distinct oceanographic hotspots. Progress in Oceanography, 219. 10.1016/j.pocean.2023.103135

57. Pease, J. B., Haak, D. C., Hahn, M. W., & Moyle, L. C. (2016). Phylogenomics Reveals Three Sources of Adaptive Variation during a Rapid Radiation. PLoS Biology, 14(2). 10.1371/journal.pbio.1002379

58. Pease, J. B., & Rosenzweig, B. K. (2018). Encoding Data Using Biological Principles: The Multisample Variant Format for Phylogenomics and Population Genomics. IEEE/ACM Transactions on Computational Biology and Bioinformatics, 15(4), 1231–1238. 10.1109/TCBB.2015.2509997

59. Richardson, D., Marancik, K., Guyon, J., Lutcavage, M., Galuardi, B., Hin, C., Walsh, H., Wildes, S., Yates, D., & Hare, J. (2016). Multiple lines of evidence for size-structured spawning migrations in western Atlantic bluefin tuna. Proceedings of the National Academy of Sciences, 113, 4262–4263. 10.1073/pnas.1607666113

60. Rooker, J., Secor, D., De Metrio, G., Schloesser, R., Block, B., & Neilson, J. (2008). Natal Homing and Connectivity in Atlantic Bluefin Tuna Populations. Science, 322, 742–744. 10.1126/science.1161473

61. Shao, F., Han, M., & Peng, Z. (2019). Evolution and diversity of transposable elements in fish genomes. Scientific Reports, 9(1). 10.1038/s41598-019-51888-1

62. Shiels, H. A., Di Maio, A., Thompson, S., & Block, B. A. (2011). Warm fish with cold hearts: Thermal plasticity of excitation-contraction coupling in bluefin tuna. Proceedings of the Royal Society B: Biological Sciences, 278(1702), 18–27. 10.1098/rspb.2010.1274

63. Shimose, T., & Farley, J. H. (2015). Age, Growth and Reproductive Biology of Bluefin Tunas.

64. Sim, S. B., Corpuz, R. L., Simmonds, T. J., & Geib, S. M. (2022). HiFiAdapterFilt, a memory efficient read processing pipeline, prevents occurrence of adapter sequence in PacBio HiFi reads and their negative impacts on genome assembly. BMC Genomics, 23(1). 10.1186/s12864-022-08375-1

65. Stanke, M., Keller, O., Gunduz, I., Hayes, A., Waack, S., & Morgenstern, B. (2006). AUGUSTUS: A b initio prediction of alternative transcripts. Nucleic Acids Research, 34(WEB. SERV. ISS.). 10.1093/nar/gkl200

66. Suda, A., Nishiki, I., Iwasaki, Y., Matsuura, A., Akita, T., Suzuki, N., & Fujiwara, A. (2019). Improvement of the Pacific bluefin tuna (Thunnus orientalis) reference genome and development of male-specific DNA markers. Scientific Reports, 9(1). 10.1038/s41598-019-50978-4

67. Supple, M. A., & Shapiro, B. (2018). Conservation of biodiversity in the genomics era. Genome Biology, 19(1). 10.1186/s13059-018-1520-3

68. Tarailo-Graovac, M., & Chen, N. (2009). Using RepeatMasker to identify repetitive elements in genomic sequences. In Current Protocols in Bioinformatics (Issue SUPPL. 25). 10.1002/0471250953.bi0410s25

69. Uchino, T., Hosoda, E., Nakamura, Y., Yasuike, M., Mekuchi, M., Sekino, M., Fujiwara, A., Sugaya, T., Tanaka, Y., Kumon, K., Agawa, Y., Sawada, Y., Sano, M., & Sakamoto, T. (2018). Genotyping-by-sequencing for construction of a new genetic linkage map and QTL analysis of growth-related traits in Pacific bluefin tuna. Aquaculture Research, 49(3), 1293–1301. 10.1111/are.13584

70. Vollger, M. R., Logsdon, G. A., Audano, P. A., Sulovari, A., Porubsky, D., Peluso, P., Wenger, A. M., Concepcion, G. T., Kronenberg, Z. N., Munson, K. M., Baker, C., Sanders, A. D., Spierings, D. C. J., Lansdorp, P. M., Surti, U., Hunkapiller, M. W., & Eichler, E. E. (2020). Improved assembly and variant detection of a haploid human genome using single-molecule, high-fidelity long reads. Annals of Human Genetics, 84(2), 125–140. 10.1111/ahg.12364

71. Walli, A., Teo, S. L. H., Boustany, A., Farwell, C. J., Williams, T., Dewar, H., Prince, E., & Block, B. A. (2009). Seasonal movements, aggregations and diving behavior of Atlantic bluefin tuna (Thunnus thynnus) revealed with archival tags. PLoS ONE, 4(7). 10.1371/journal.pone.0006151

72. Watanabe, Y. Y., Goldman, K. J., Caselle, J. E., Chapman, D. D., & Papastamatiou, Y. P. (2015). Comparative analyses of animal-tracking data reveal ecological significance of endothermy in fishes. Proceedings of the National Academy of Sciences of the United States of America, 112(19), 6104–6109. 10.1073/pnas.1500316112

73. Wenger, A. M., Peluso, P., Rowell, W. J., Chang, P. C., Hall, R. J., Concepcion, G. T., Ebler, J., Fungtammasan, A., Kolesnikov, A., Olson, N. D., Töpfer, A., Alonge, M., Mahmoud, M., Qian, Y., Chin, C. S., Phillippy, A. M., Schatz, M. C., Myers, G., DePristo, M. A., … Hunkapiller, M. W. (2019). Accurate circular consensus long-read sequencing improves variant detection and assembly of a human genome. Nature Biotechnology, 37(10), 1155–1162. 10.1038/s41587-019-0217-9

74. Whitlock, R. E., Hazen, E. L., Walli, A., Farwell, C., Bograd, S. J., Foley, D. G., Castleton, M., & Block, B. A. (2015). Direct quantification of energy intake in an apex marine predator suggests physiology is a key driver of migrations. Science Advances, 1(8). 10.1126/sciadv.1400270

75. Yang, Z. (2007). PAML 4: Phylogenetic analysis by maximum likelihood. Molecular Biology and Evolution, 24(8), 1586–1591. 10.1093/molbev/msm088

76. Zhao, X., Huang, Y., Bian, C., You, X., Zhang, X., Chen, J., Wang, M., Hu, C., Xu, Y., Xu, J., & Shi, Q. (2022). Whole genome sequencing of the fast-swimming Southern bluefin tuna (Thunnus maccoyii). Frontiers in Genetics, 13. 10.3389/fgene.2022.1020017

77. Zook, J. M., Catoe, D., McDaniel, J., Vang, L., Spies, N., Sidow, A., Weng, Z., Liu, Y., Mason, C. E., Alexander, N., Henaff, E., McIntyre, A. B. R., Chandramohan, D., Chen, F., Jaeger, E., Moshrefi, A., Pham, K., Stedman, W., Liang, T., … Salit, M. (2016). Extensive sequencing of seven human genomes to characterize benchmark reference materials. Scientific Data, 3. 10.1038/sdata.2016.25

78. Zook, J. M., McDaniel, J., Olson, N. D., Wagner, J., Parikh, H., Heaton, H., Irvine, S. A., Trigg, L., Truty, R., McLean, C. Y., De La Vega, F. M., Xiao, C., Sherry, S., & Salit, M. (2019). An open resource for accurately benchmarking small variant and reference calls. Nature Biotechnology, 37(5), 561–566. 10.1038/s41587-019-0074-6

